# Characterising time-on-task effects on oscillatory and aperiodic EEG components and their co-variation with visual task performance

**DOI:** 10.1101/2024.04.19.590227

**Authors:** Martina Kopčanová, Gregor Thut, Christopher SY Benwell, Christian Keitel

## Abstract

Research on brain-behaviour relationships often makes the implicit assumption that these derive from a co-variation of stochastic fluctuations in brain activity and performance across trials of an experiment. However, challenging this assumption, oscillatory brain activity, as well as indicators of performance, such as response speed, can show systematic trends with time on task. Here we tested whether time-on-task trends explain a range of relationships between oscillatory brain activity and response speed, accuracy as well as decision confidence. Thirty-six participants performed 900 trials of a two-alternative forced choice visual discrimination task with confidence ratings. Pre- and post-stimulus spectral power (1-40Hz) and aperiodic (i.e., non-oscillatory) components were compared across blocks of the experimental session and tested for relationships with behavioural performance. We found that time-on-task effects on oscillatory EEG activity were primarily localised within the alpha band, with alpha power increasing and peak alpha frequency decreasing over time, even when controlling for aperiodic contributions. Aperiodic, broadband activity on the other hand did not show time-on-task effects in our data set. Importantly, time-on-task effects in alpha frequency and power explained variability in single-trial reaction times, and controlling for time-on-task effectively removed these relationships. However, time-on-task effects did not affect other EEG signatures of behavioural performance, including post-stimulus predictors of single-trial decision confidence. Our results dissociate alpha-band brain-behaviour relationships that can be explained away by time-on-task from those that remain after accounting for it - thereby further specifying the potential functional roles of alpha in human visual perception.

## Introduction

Neural oscillations arising from fluctuations in synchronised networks represent a fundamental characteristic of the brain across species (Buzsáki et al., 2013). This oscillatory activity, measurable with EEG, has repeatedly been linked to cognitive functions and behaviour in various domains (Başar et al., 2001; Fries, 2015; Thut et al., 2012; Ward, 2003). Hence, neural oscillations have been proposed to play a crucial role in global functions such as encoding and transfer of information between large-scale neural networks (Fries, 2015; Keitel et al., 2022). For instance, fluctuations in both spontaneous (pre-stimulus) and stimulus-related changes in alpha band activity (8-13Hz) have been linked to numerous behavioural outcomes in e.g. perception (Babiloni et al., 2006; Benwell et al., 2022; Busch et al., 2009; Chaumon & Busch, 2014; Craddock et al., 2017) and memory (Hanslmayr et al., 2012; Klimesch et al., 2006), and have been assigned different functional roles, such as functional inhibition (Foxe & Snyder, 2011) and information gating (Jensen & Mazaheri, 2010).

In addition to spontaneous and state-/stimulus-dependent fluctuations, oscillations exhibit systematic non-stationarities on shorter timescales, in the order of seconds (Cohen, 2014) and across the lengths of standard EEG experiments (>1h, Benwell et al., 2019) that may be of functional importance. For instance, Benwell et al. (2019) showed evidence for partially independent drifts in alpha power and peak frequency over the course of an ∼1h experimental session, in line with previous studies examining the effects of sustained task performance on EEG activity (Boksem et al., 2005; Stoll et al., 2016). These findings highlight that changes in alpha oscillations cannot be assumed to be purely stochastic (i.e., due to trial-by-trial variability) but are rather partially deterministic, implying sources of variance that act on longer time scales of several seconds to hours, e.g., vigilance decrements (Hemmerich et al., 2023; Martel et al., 2014; Pershin et al., 2023) or fluctuations in arousal. Separating the relative influences of stochastic versus deterministic sources of alpha variability on perception and behaviour is thus crucial for understanding its functional role(s). Importantly, in cases where the behaviour of interest also shows non-stationarity across the experiment (e.g., Benwell et al., 2018; Doll et al., 2015; Fründ et al., 2011), any brain-behaviour relationships observed may be confounded, arising due to time-on-task effects on both variables. Without accounting for time-on-task, such effects may be misinterpreted to represent functional relationships (Benwell et al., 2018; Schaworonkow et al., 2015). Alternatively, systematic changes in alpha activity over time may represent a neural mechanism governing systematic changes in behaviour.

It remains unclear whether EEG time-on-task effects extend beyond the alpha band (though see Arnau et al., 2021) and to what extent they differ depending on the task performed. Previous research examining time-on-task effects on EEG activity has not considered the broadband aperiodic component, which is superimposed with oscillatory activity in the power spectrum (Donoghue, Haller, et al., 2020). Aperiodic activity often confounds estimates of band-specific spectral power (Donoghue, Dominguez, et al., 2020; Kopčanová et al., 2024) raising the possibility that previously observed alpha-band time-on-task effects can be explained by changes in the aperiodic component. Moreover, aperiodic activity itself has been proposed to index neural excitation/inhibition balance (Gao et al., 2017), and, accordingly, has been found to co-vary with cognitive performance (Ouyang et al., 2020; Waschke et al., 2021).

Here we examined time-on-task related fluctuations in behaviour and oscillatory activity while participants performed a 2-alternative forced choice visual discrimination task with confidence ratings. This allowed us to investigate the presence of any systematic non-stationarities in accuracy (type-1), confidence (type-2 decisions) and reaction times as well as their relationship to spectral EEG measures. We tested for time-on-task effects across a wide 1-40Hz frequency spectrum and parameterized the EEG spectra to determine whether any time-on-task effects were driven by periodic or aperiodic changes. Single-trial multiple regression analyses were used to test whether time-on-task effects confounded any observed brain-behaviour relationships.

## Methods

### Participants

Thirty-seven participants completed a visual 2-AFC discrimination task while their EEG was recorded. Participants were included if they were 18-40 years old, had normal or corrected-to-normal vision, no colour-blindness, and no history of neurological disorders. After task performance checks (see Behavioural analysis), 36 participants were included in the analysis (27 female, 33 right-handed, age: *M* = 22.9, *SD* = 4.4, range = 18-35). They reported sleeping 7.3 hours on average prior to testing, and 17 wore glasses or contact lenses. We obtained written informed consent from all participants according to the Declaration of Helsinki. All participants were financially compensated for their time (£20). The School of Social Sciences Ethics Committee and the University of Dundee approved the study (UoD-SHSL-PSY-TPG-2022-220).

### Task and experimental design

The study involved a 2-alternative forced choice (2-AFC) discrimination task (adapted from Benwell et al., 2022) with confidence and attentional ratings, as well as simultaneous 32-channel EEG recording. Participants were required to discriminate numbers from letters and report confidence in their decisions (see Figure 1). On a subset of trials, they additionally rated their attention to the task on a continuous scale (1 to 10) – however, this data will not be used for the present analysis.

**Figure 1.**
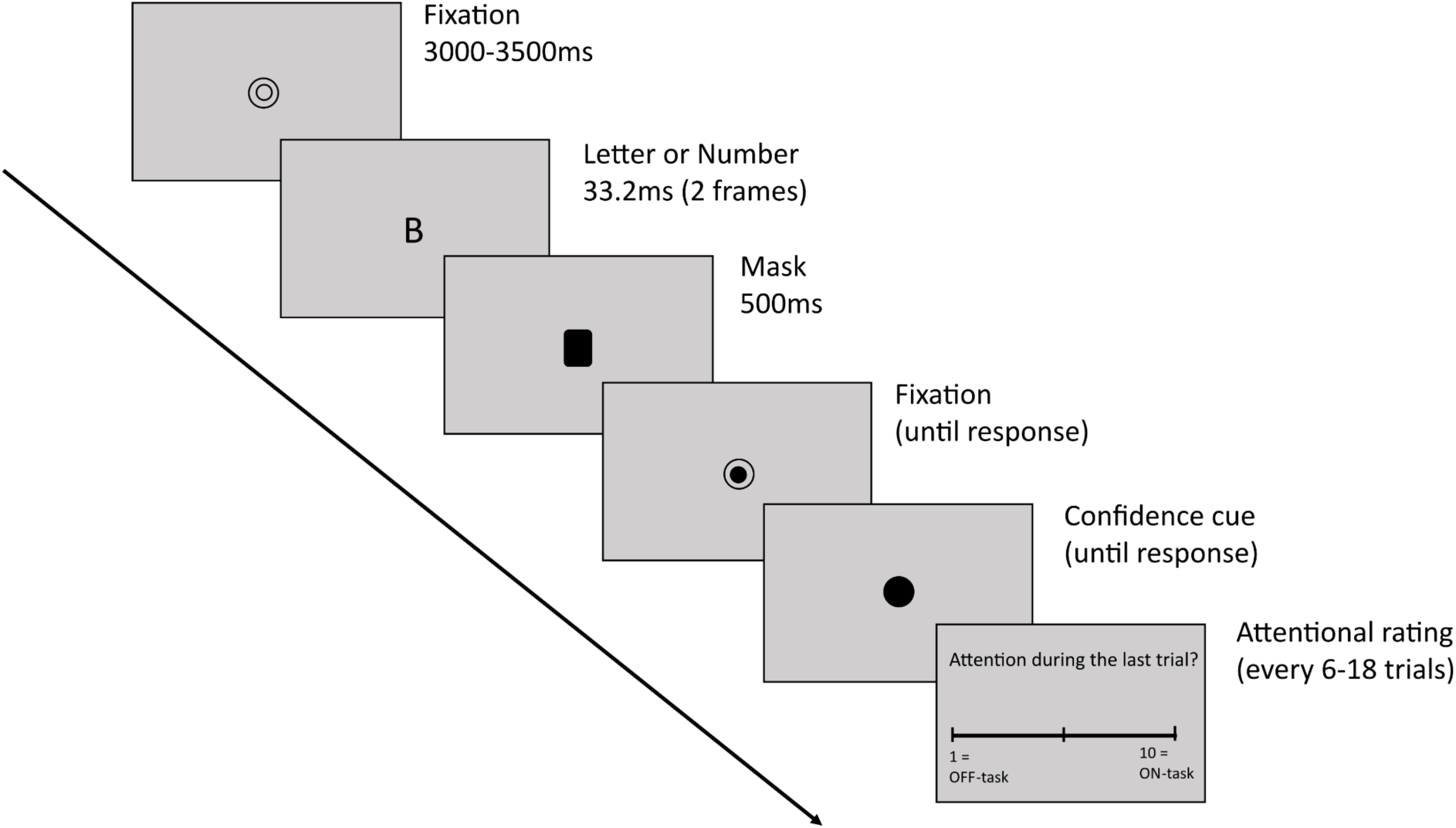
Task design. The task began with a fixation circle presented for a varying duration of 3000-3500ms on a gray background. It was followed by the stimulus (either a number or a letter) presented for ∼33ms and quickly masked with a black rectangle for 500ms. Participants could respond immediately. A half-filled fixation circle remained on the screen until they responded and was followed by a black filled circle indicating that a confidence rating should now be given. On a subset of trials participants also rated their attention on a continuous scale from 1 to 10.

We used the PsychoPy software (Peirce et al., 2019) to present the stimuli on a 53×30cm monitor with 1920×1080 resolution and 60Hz refresh rate. Recordings took place in a sound-attenuated booth in our local EEG laboratory. Participants sat 57cm from the screen and the room was dimly lit. Each trial begun with a black fixation circle (1×1 cm) on a gray background presented for a duration randomly varying between 3000-3500ms. The fixation was followed by the stimulus (2×1.3 cm), either a letter randomly selected from 18 consonants (B, C, D, F, G, H, J, K, L, M, N, P, R, S, T, V, X, Z) or a number selected from digits 1-9. Numbers or letters presentations were equiprobable. The stimuli were briefly presented for 2 frames (∼= 33.3ms), and immediately masked with an overlay of all possible letters and numbers (duration = 500ms, 2×1.3 cm). Difficulty was manipulated by changing the contrast of the stimuli to 5 difficulty levels, selected prior to the experiment. We created five difficulty levels by varying the opacity of the stimuli in PsychoPy to the following levels (hardest to easiest): 0.075, 0.15, 0.225, 0.3, 0.375. This range of stimulus contrast levels was selected after running a behavioural pilot with 15 levels (0.05-0.4, in increments of 0.025) and selecting the range with optimal psychometric functions. The mask was followed by a response-cue fixation circle (half-filled circle) that stayed on the screen until the participants made their 1^st^-order response. They were required to use their left hand to press ‘a’ if they saw a number, or ‘s’ if they saw a letter. Afterwards, a confidence-cue fixation appeared (fully filled circle) and remained on the screen until participants rated their confidence. To do so, with their right hand, participants pressed ‘k’ for confident in the choice being correct or ‘l’ for not confident. There were 900 trials in total, split into 5 blocks of 180 trials and the trial order was fully randomised within blocks. On 15 trials within each block (75 trials total), an attentional rating scale appeared after the participants rated their confidence. Participants were asked to what extent they were focused *on* or *off* the task during the preceding trial on a scale of 1 to 10 (1 = complete *off* task attention, 10 = complete *on* task attention). The attentional probes were spaced every 6-18 trials (with delays sampled from a uniform distribution, mean delay 11 trials). To familiarise themselves with the task, all participants completed 20 practice trials at the beginning of the experiment with feedback. Overall, the experiment took ∼2-2.5 hours including the EEG set-up. Participants took self-paced breaks between blocks.

### Behavioural analysis

We evaluated the effectiveness of the task difficulty manipulation by plotting the psychometric functions for both accuracy and confidence across all stimulus contrast levels. One participant was excluded from the study at this stage due to chance performance throughout the task (mean proportion correct = 50%). Additionally, we tested for time-on-task effects on behaviour by calculating the accuracy (type-1), confidence (type-2), and response RT psychometric functions for each of the five blocks separately. To statistically test for main effects of time-on-task (i.e., block) and stimulus contrast (i.e., difficulty) and their interaction, we used 5×5 within-participant ANOVAs with block number and stimulus contrast level as the independent variables and proportion correct/confident, or mean RT, as the dependent variables. One-way ANOVAs with corrections for multiple comparisons (Tukey HSD) were used to follow up any significant main and/or interaction effects. We applied the Greenhouse-Geisser correction (*p_gg_*) to any models where the Mauchly’s test of sphericity indicated unequal variances (p < .05).

### EEG acquisition and pre-processing

We recorded continuous EEG with a 32-channel ActiveTwo system (Biosemi, The Netherlands) at 512Hz sampling rate. We placed the scalp electrodes according to the 10-20 International system (Oostenveld & Praamstra, 2001) and four electrooculographic electrodes were additionally placed at the outer canthi of each eye and above and below the right eye.

We pre-processed the EEG data offline with custom-written scripts in MATLAB (R2022a) (Mathworks, USA), using Fieldtrip (Oostenveld et al., 2011) and EEGLAB (Delorme & Makeig, 2004). A zero-phase 2nd-order Butterworth filter was used to apply low-pass (100Hz) and high-pass (0.5Hz) filters to the data. We then obtained 4-second epochs spanning from -2.5 to 1.5s relative to stimulus onset on every trial. Excessively noisy channels were removed without interpolation after visual inspection (sample mean = 0.22 removed channels, range = 0-3). Next, we average-referenced the data and removed excessively noisy trials with a semi-automated artifact rejection procedure. Noisy trials were selected based on 1) extreme amplitudes ( ± 75 µV), 2) joint probability (‘pop_jointprob’ (Delorme & Makeig, 2004), with 3.5SD single-channel limit and 3SD for global limit), 3) kurtosis (local threshold 5SD, global threshold 3SD, ‘pop_rejkurt’ (Delorme & Makeig, 2004)), and 4) visual inspection. We then used the ‘runica’ function in EEGLAB (Delorme & Makeig, 2004) to run an independent component analysis (ICA) using the default settings. Afterwards, we used the Multiple Artifact Rejection Algorithm (MARA) (Winkler et al., 2011) to semi-automatically remove the components corresponding to ocular and muscle activity, or transient channel noise. Any remaining noisy epochs were then removed resulting in retaining 870 of 900 trials per participant on average (range = 824-897). Lastly, we interpolated any removed channels with a spherical spline method to allow group analysis and plotting topographies.

### EEG spectral decomposition

To obtain single-trial power spectra for both the pre-stimulus (-1 to 0s) and post-stimulus time (0 to 1s) periods separately, we used an FFT applied to the Hanning-tapered data using the ‘mtmfft’ function from Fieldtrip (Oostenveld et al., 2011). Prior to the FFT data were zero-padded to a length of 10s to achieve a 0.1Hz frequency resolution across the range of 1-40Hz.

### Spectral Parameterization

Single-trial power spectra were then averaged across all 32 electrodes (separately for pre- and post-stimulus data) and used for spectral parameterization. We used the ‘specparam’ toolbox (Donoghue, Haller, et al., 2020) to isolate the periodic and aperiodic activity in the power spectra. Parametrization was applied to the frequency range of 3-40Hz, using the ‘fixed’ mode, with a peak detection threshold of 2 SD, peak width limits set to 2-15Hz, and a maximum number of to-be-detected peaks of 4, similar to previous studies (Donoghue, Dominguez, et al., 2020; Donoghue, Haller, et al., 2020; Robertson et al., 2019). The single trial fits were very good overall for both pre-stimulus spectra (grand mean R^2^ (error) = 0.924 (0.139)), with a range of R^2^ = 0.645-0.985 across participants, and post-stimulus spectra (grand mean R^2^(error) = 0.923 (0.142), with a range of R^2^ = 0.655-0.984). The single-trial estimates of aperiodic exponents were then extracted and used in subsequent statistical analyses.

The parameterized spectra were also used to identify single-trial oscillatory peaks falling into an extended alpha range (6-15Hz). Where an alpha peak was detected (pre-stimulus: mean = 823 trials, range = 671-889; post-stimulus: mean = 784 trials, range = 546-877), we extracted its center frequency (calculated by ‘specparam’) and aperiodic-adjusted power.

### Time-frequency analysis

We also performed a Fourier-based time-frequency decomposition on the artefact-free single-trial data for each electrode using the ‘ft_freqanalysis’ function and ‘mtmconvol’ method (Fieldtrip, Oostenveld et al., 2011). We maintained temporal resolution by decomposing overlapping 0.5-second sections of time-series data that were consecutively shifted forward by 0.02 s. We Hanning tapered the data segments and zero-padded to a length of 1 second to achieve a frequency resolution of 1 Hz over a range of 1-40Hz. To obtain single-trial spectral power values, we squared the absolute values of the complex-valued time-frequency estimates. Data epochs from -1 to 1s relative to stimulus onset were then used in subsequent statistical analyses.

### Statistical analyses

We performed all statistics in MATLAB 2022a (Mathworks, USA) using the implementations of standard procedures (ANOVA, regression) where applicable, in combination with custom-written code for data preparation.

#### Time-on-task effects on EEG power spectra

To investigate how EEG power spectra changed with time-on-task we adopted a non-parametric cluster-based permutation approach (Maris & Oostenveld, 2007). Specifically, we tested whether the power spectra differed significantly across the five experimental blocks. This analysis was performed separately for pre- and post-stimulus power spectra.

First, for each participant, we computed the mean amplitude at each channel-frequency point separately for each block. To test for systematic group-level effects, we then combined the average amplitudes across all participants and compared them with a dependent-samples F-test (‘ft_freqstatistics’, method: ‘montecarlo’ in Fieldtrip (Oostenveld et al., 2011)). Based on the initial F-tests, clusters were formed by grouping all neighbouring significant F-values (*p* <.05). Clusters needed to consist of at least two neighbouring electrodes. Spatially, we defined the neighbours based on the ‘*biosemi32-neighb.mat’* template from Fieldtrip (Oostenveld et al., 2011), with a mean of 6.0 neighbours per electrode. The significance of the summed F-values (i.e., cluster statistics) was determined against a data-driven null-distribution obtained by repeating the whole procedure 2000 times while randomly permuting the data. The cluster-level significance threshold was *p* < .05.

To test the relationships between time-on-task and the alpha peak power, frequency, and the aperiodic exponent, we averaged the single-trial estimates of each measure for each block. Then, a within-participant ANOVA with block as independent variable and EEG measure as dependent variable was used to test for time-on-task effects. Again, this analysis was performed separately for both pre- and post-stimulus EEG measure estimates.

#### Testing whether time-on-task accounts for EEG-behaviour relationships

To investigate whether any time-on-task effects on EEG activity affected brain-behaviour relationships we performed a single-trial multiple regression analysis (for similar approach see Benwell et al., 2017, 2022; Kopčanová et al., 2023). We used a hierarchical two-stage estimation (Friston, 2008) to incorporate participant level variability into group level statistics.

*Within-participant* relationships between spectral (time-frequency) power and each of the behavioural variables (accuracy, confidence, response RT), as well as trial-order and stimulus contrast level were tested with a multiple regression for each time-frequency-electrode combination. This allowed us to statistically separate the unique effects of time-on-task and behaviour on the EEG outcome. The regression model was applied using the ‘fitlm’ function using the least-squares criterion (Mathworks, USA) and was defined as follows:

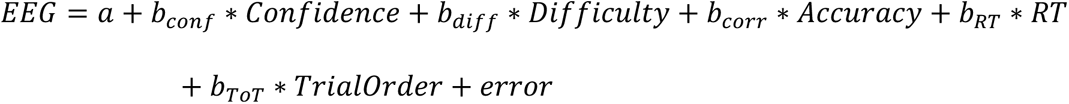

We thus obtained the regression coefficients (*b_conf_*, *b_corr_*, *b_diff_*, *b_RT_*, *b_ToT_*) for the EEG-behaviour/time-on-task relationships that represented the strength and direction of the association between spectral power and each predictor while the other predictors were controlled for. All regression models were computed without trial-order first, and then trail-order was added as an additional predictor in a separate model to test whether accounting for trial order would abolish any observed brain-behaviour relationships. Additionally, predictive performance of both models was compared by first evaluating the subject-level Akaike Information Criterion (Cavanaugh & Neath, 2019), then contrasting resulting AIC values by means of a paired t-test (two-tailed).

*Group-level* statistics were computed using cluster-based permutation testing of the obtained regression coefficients against zero (‘ft_freqstatistics’, two-tailed dependent-samples t-test, method: ‘montecarlo’ in Fieldtrip (Oostenveld et al., 2011)). The procedure was similar to that described above for EEG power spectra obtained from the FFT analysis with a cluster p-value threshold = .025.

We used this multiple regression approach to examine whether any time-on-task effects influenced the relationships between peak alpha power/frequency and behaviour as well.

### Data and code availability

Data and code are openly available online. The data can be accessed through OSF (<u>here</u>), and the code can be found on GitHub (<u>here</u>).

## Results

### Behavioural results

We first evaluated the effectiveness of the stimulus contrast manipulation and time-on-task effects on three behavioural measures: accuracy (type-1 performance), confidence (type-2 performance), and response RTs. Figure 2 shows the psychometric functions for each behaviour type, plotted separately for each task block.

**Figure 2:**
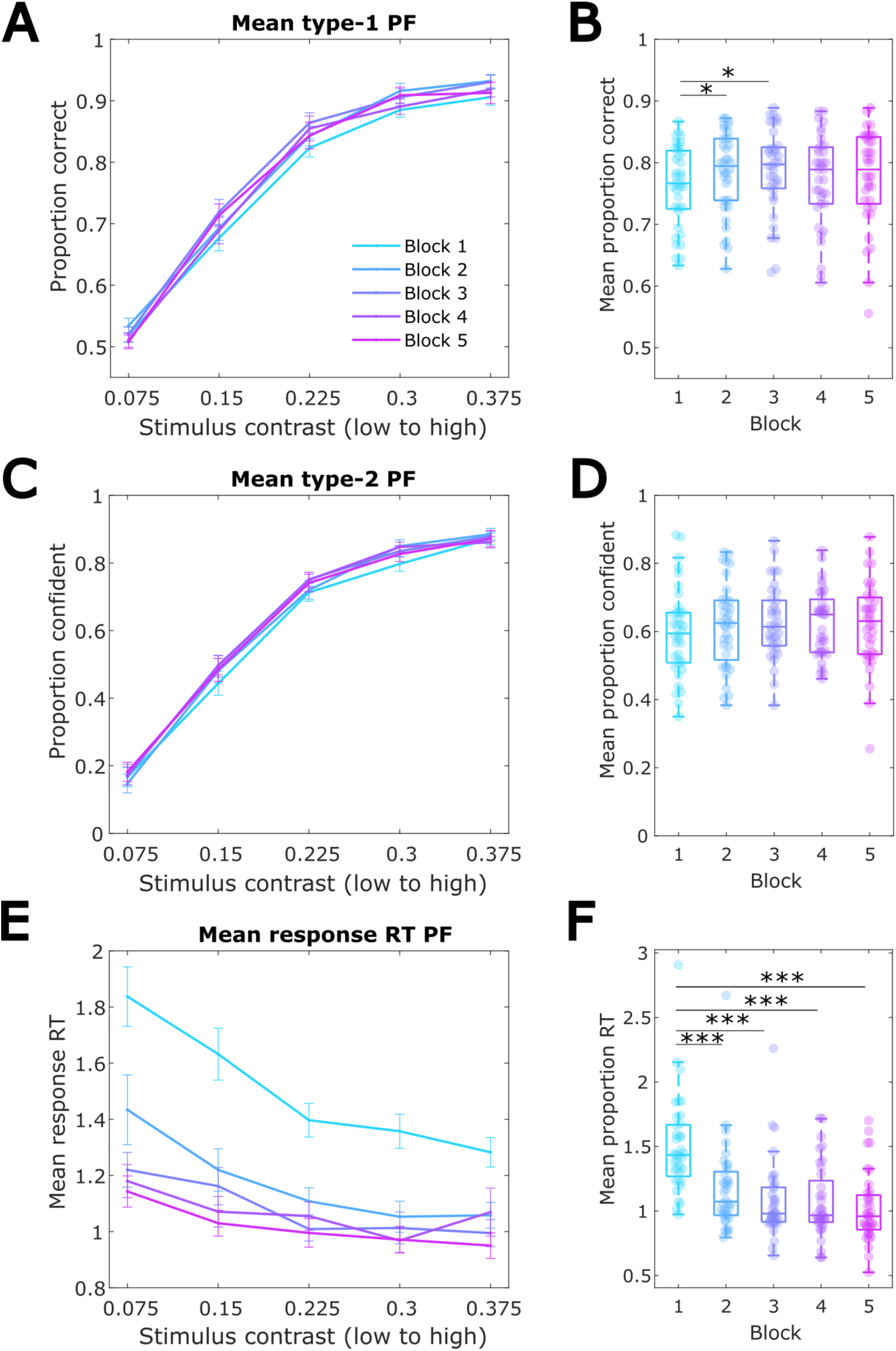
Time-on-task and stimulus contrast effects on behaviour. (A) Mean proportion of correct responses per stimulus contrast level within each of the five blocks. Error bars denote the between-participant standard error of the mean. (B) Mean proportion correct responses per block. (C) Mean proportion of confident responses per stimulus contrast level in each block. (D) Mean proportion of confident responses per block (E) Mean response RTs per stimulus contrast level and block. (F) Mean RTs per block. * *p* < .05 ** *p* < .001 *** *p* < .0001

A 5-by-5 ANOVA showed that accuracy increased with higher stimulus contrast as expected: F(1.49,52.1) = 494.761, *p_gg_* < .0001, η_p_^2^ = .934 (see also Figure 2A). The main effect of time-on-task on accuracy was also significant, F(1.49,52.1) = 3.703, *p_gg_* = .018, η_p_^2^ = .096, whilst the interaction between block and stimulus contrast level was not significant, F(5.95,208.4) = 1.212, *p_gg_* = .284. A follow-up one-way ANOVA with multiple comparison correction testing the effect of block on accuracy across all evidence levels (F(2.67,93.31) = 3.703, *p_gg_* = .018, η^2^ = .097) showed that the proportion of correct responses increased at the beginning of the task from block 1 to block 2 (p = .016) and from block 1 to block 3 (p = .0008), but no other comparisons were significant (Figure 2B).

Like accuracy (type-1 performance), the proportion of confident responses also increased as a function of stimulus contrast, F(0.88,30.63) = 372.427, *p_gg_* <.0001, η_p_^2^ = .914 (Figure 2C). However, there was no significant main effect of time-on-task on confidence, F(0.88,30.63) = 1.291, *p_gg_* = .282 (Figure 2D). The effect of stimulus contrast on confidence did not vary across blocks, F(3.5,122.52) = 1.766, *p_gg_* = .071.

As Figure 2E shows, participants’ responses got faster with increasing stimulus contrast, F(0.68,23.81) = 18.590, *p_gg_* <.0001, η_p_^2^ = .347, as well as with time-on-task, F(0.68,23.81) = 35.690, *p_gg_* <.0001, η_p_^2^ = .505. A follow up one-way ANOVA with multiple comparisons of the main time-on-task effect on RTs (F(2.82,98.68) = 35.673, *p_gg_* <. 0001, η^2^ = .505) showed that RTs were significantly slower in block 1 compared to blocks 2, 3, 4, and 5 (p’s <.0001) and no other comparisons were significant (see also Figure 2F). Additionally, there was a significant interaction between block and stimulus contrast on RTs, F(2.72,95.22) = 4.696, *p_gg_* =.0006, η_p_^2^ = .118. Follow up one-way ANOVAs with stimulus contrast as independent variable and RT as dependent variable carried out separately at each block showed that this interaction is driven by differences in the effect of the stimulus contrast manipulation on RTs across blocks. Specifically, as Table S1 the Supplementary Material shows, block 4 did not show the main effect of stimulus contrast on RTs. No differences in the effect of block on RTs across the stimulus contrast levels were found (see Supplementary Table S2).

In summary, the task difficulty manipulation was effective. Importantly, the improvement in accuracy and speed-up of RTs, primarily shown in comparisons between the first and later blocks, may indicate practice effects on behaviour. Confidence ratings did not show systematic variability between blocks.

### EEG results

#### Time-on-task effects on spectral power are strongest in the alpha band

We next tested time-on-task effects on the EEG power spectra across the 1-40Hz range. Figure 3A-B plots the full scalp mean spectra for each task block separately for 1-sec pre- and post-stimulus periods. To statistically test for any significant differences in power spectra across all electrodes we used a cluster-based permutation F-test. We found significant clusters in alpha frequencies in both the pre-stimulus, F_sum_ = 13,192, p_clust_ = .0005 (3.7-11.9Hz), and the post-stimulus, F_sum_ = 11,188, p_clust_ =.001 (5.2-12.4Hz), periods. Additionally, there were significant clusters in beta frequencies, with an F_sum_ = 1,684 and p_clust_ = .0435 (12.7-18.7Hz) in the pre-stimulus period, and with an F_sum_ = 2,711 and p_clust_ = .024 (13-18.9Hz) in the post-stimulus period. The topographies in Figure 3C-D show that the grand-average spectral power in both the alpha and beta bands showed similar topographical distributions between the first and the last blocks, but the overall amplitudes increased with time. The alpha time-on-task effect was significant across the whole scalp in both pre- and post-stimulus periods, whilst the beta effect was constrained to frontal electrodes in the pre-stimulus period, and to frontal and parieto-occipital electrodes in the post-stimulus period.

**Figure 3.**
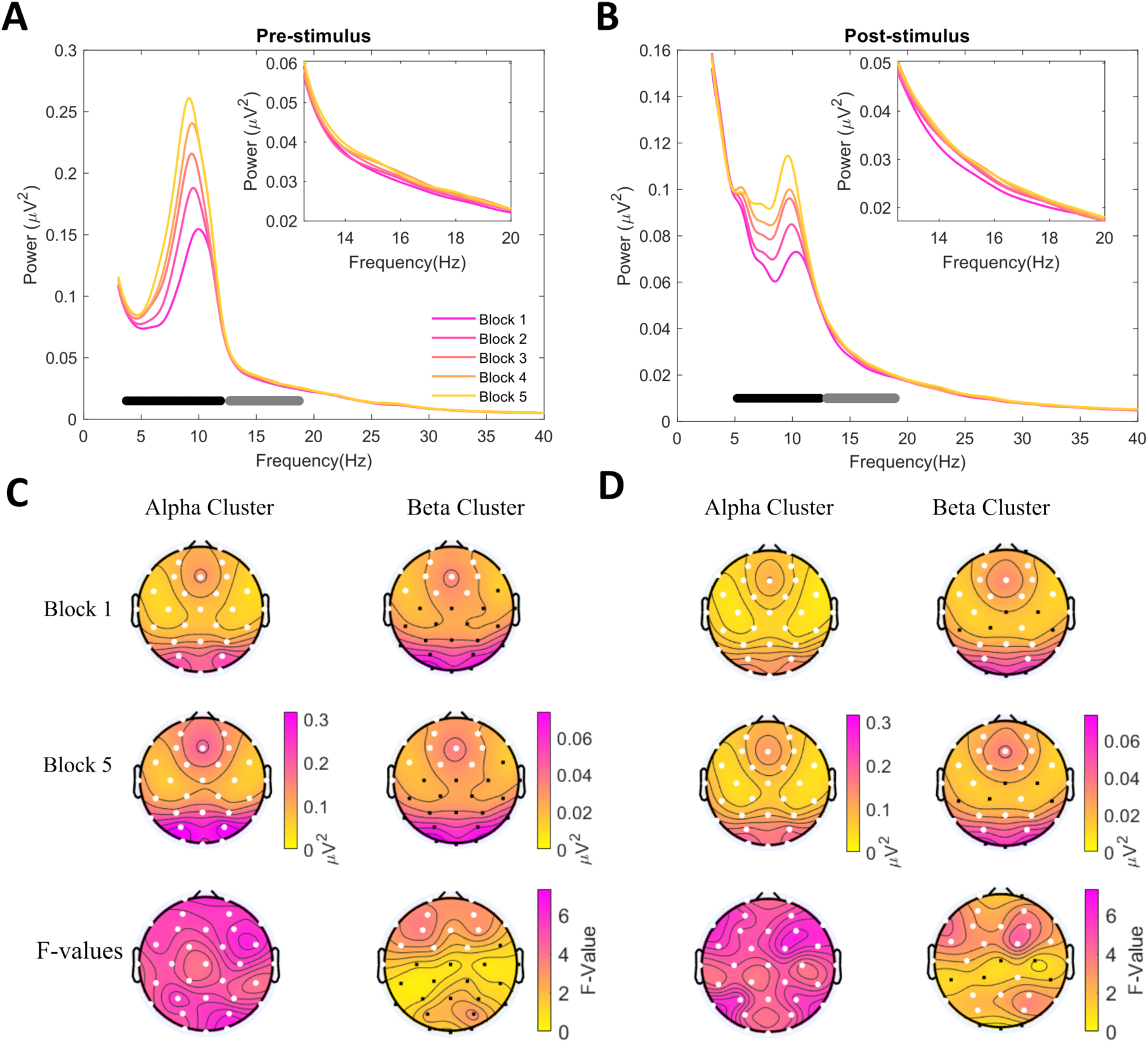
Time-on-task effect on 1-40Hz power spectra across blocks. (A-B) Scalp mean grand-average power spectra plotted separately for each experimental block. The black and gray horizontal lines indicate the frequency ranges included in the two significant clusters found when testing the differences in amplitude across all 5 blocks with a dependent-samples F-test (α = .05). The top inset plots the power spectra in the beta frequencies. (A) Pre-stimulus power and (B) post-stimulus power. (C-D) Topographies of mean raw power in block 1 (top row) and block 5 (bottom row) as well as the mean F-values. Topographies are plotted separately for alpha and beta frequencies. The electrodes in white denote the channels that were included in the significant alpha/beta clusters.

Figure 4 further plots the within-block changes in alpha and beta power across the experiment. It shows that both alpha and beta band power increased linearly throughout the experiment, however, alpha power showed greater fluctuations within the blocks than beta power. Notably, alpha power was drastically reduced for the first trial bin in each block, suggesting a relative ‘resetting’ of power during breaks which quickly increased again with time-on-task. This pattern of changes was similar in the pre-stimulus (Fig. 4A) and post-stimulus (Fig. 4B) periods.

**Figure 4.**
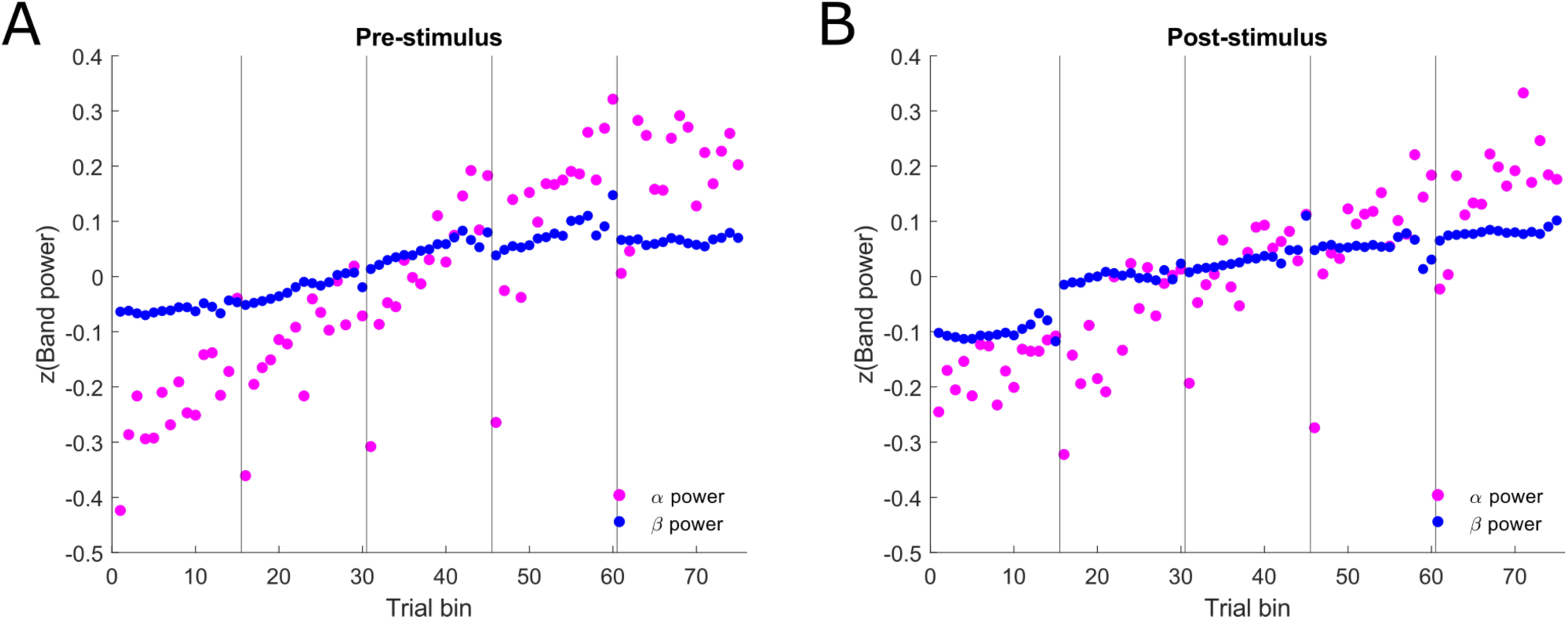
Within-block evolutions of alpha and beta band power. Mean alpha and beta power (z-scored) were calculated for each trial bin based on the significant clusters showing the effect of block on spectral power in Figure 3. Each block of trials was split into 15 bins so that the trial numbers in each bin were approximately equal. The number of trials in each bin ranged from 10-12. Vertical black lines denote block onset. **A** pre-stimulus power. **B** post-stimulus power.

#### Time-on-task changes in peak alpha frequency and power are oscillatory and not influenced by aperiodic activity

Whilst the analysis above and previous research (Benwell et al., 2019; Stoll et al., 2016) suggest power changes with time-on-task, they do not account for potential non-stationarities in the broadband aperiodic activity. Hence, focusing on the activity in the alpha band that showed the strongest time-on-task alterations, we next investigated time-on-task effects on purely oscillatory alpha activity, as parameterized by ‘specparam’ (Donoghue, Haller, et al., 2020), as well as any possible non-stationarities in the aperiodic exponent itself.

Figure 5A-B shows the full-scalp mean aperiodic-adjusted power in the alpha band (8-13Hz) for each experimental block, separately for pre- and post-stimulus windows. The vertical dashed lines additionally illustrate the mean alpha peak frequency shift across the five task blocks. As expected, Figure 5A-B illustrates that alpha power increased over blocks whilst peak alpha frequency decreased in both the pre- and post-stimulus periods. To test whether these shifts were statistically significant, we extracted the trials for which an alpha (6-15Hz) peak was identified and compared the mean center frequency and peak power (over and above the aperiodic component) across the five blocks with a one-way ANOVA. We found that alpha center frequency decreased significantly with time in both the pre-stimulus, F(4,140) = 20.752, p <.0001, η^2^ = .372 (all post-hoc pairwise comparisons p < .05 except block, 2 vs 3, and 4 vs 5), as well as post-stimulus windows, F(4,140) = 5.429, p = .0004, η^2^ = .134 (only blocks 2 and 5 differed significantly, p = .019, all other comparisons p > .05) (Figure 5C-D). In line with the aperiodic-uncorrected results above, alpha peak power also showed a significant increase in both the pre-stimulus, F(4,140) = 21.238, p < .0001, η^2^ = .378 (all post-hoc pairwise comparisons p <.05, except block 3 vs 4, 3 vs 5, and 4 vs 5), as well as the post-stimulus period, F(4,140) = 10.484, p < .0001, η^2^ = .231 (post-hoc pairwise comparisons showed block 5 differed significantly from blocks 1, 2, and 3; and block 4 differed from blocks 1 and 2) (Figure 5E-F).

**Figure 5.**
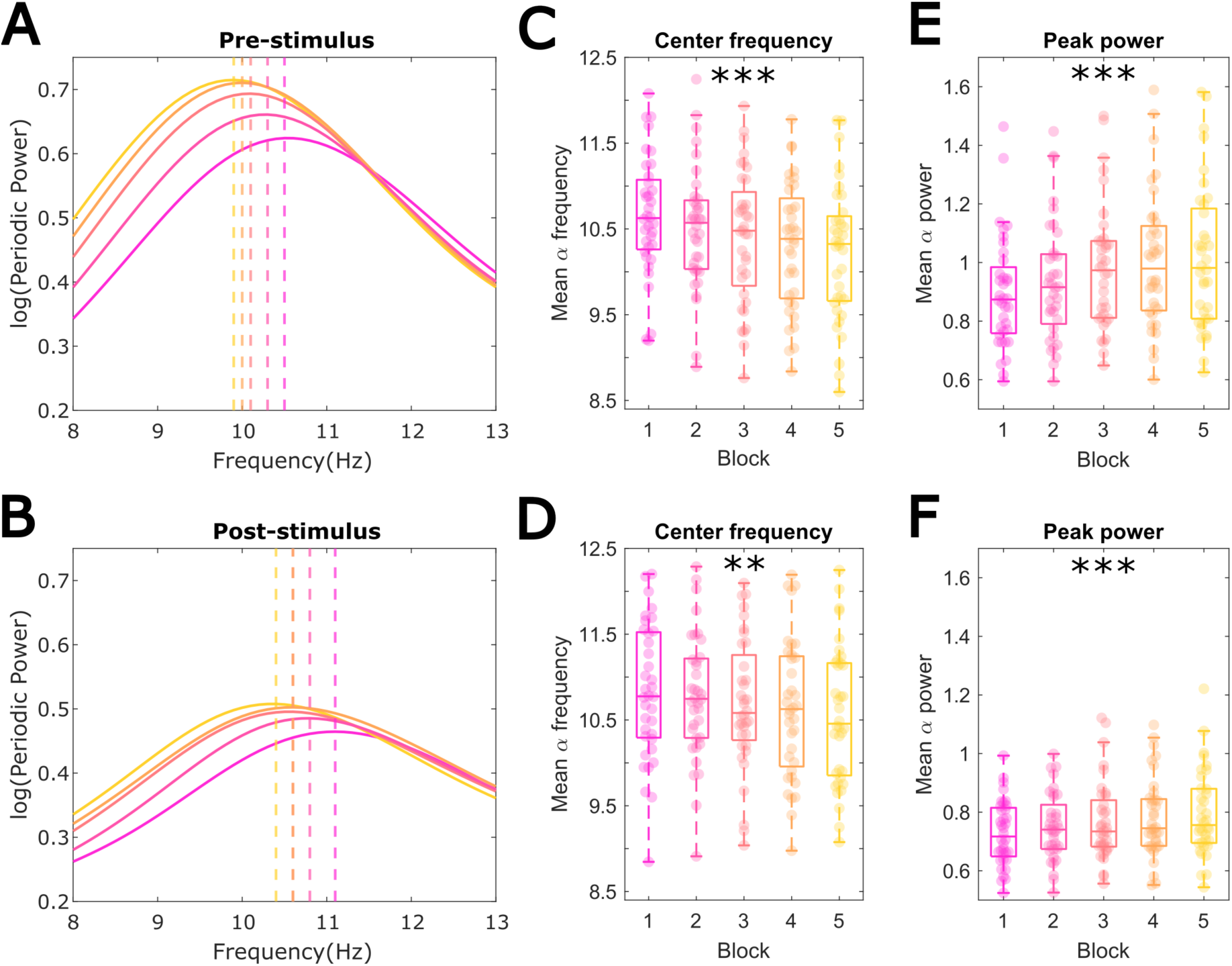
Aperiodic-adjusted alpha peak power and frequency across blocks. (A) Pre-stimulus scalp averaged mean alpha power (8-13) for each block. Vertical dashed lines denote the point at which the adjusted-alpha power is the highest. (B) Post-stimulus aperiodic-adjusted alpha power and its peak frequencies across blocks. (C) Pre-stimulus and (D) post-stimulus mean ‘specparam’ identified alpha peak center frequency per block decreased significantly with time-on-task. (E) Pre-stimulus and (F) post-stimulus mean alpha peak power increased across blocks. * p < .05 ** p < .001 *** p < .0001

Only trials with an identifiable alpha peak (identified with ‘specparam’) were included in this analysis. Overall, trials in the second half of the experiment were more likely to contain an alpha peak (on average, ∼64% of trials with no pre-stimulus alpha were from the 1^st^ half of the experiment, while ∼36% from the 2^nd^ half). While this may have slightly imbalanced the precision of the power and frequency estimates across the experiment it will not have systematically biased them.

While aimed at confirming the oscillatory nature of the alpha-band time-on-task effects, ‘specparam’ also allowed quantifying the aperiodic component of the spectrum and testing whether this showed similar effects. Figure 6A-B, shows the mean aperiodic exponent estimates for each block, plotted separately for pre- and post-stimulus data. One-way within-participant ANOVAs indicated that there were no significant time-on-task changes in either pre-stimulus, F(4,140) = 2.032, p = .093, or post-stimulus, F(4,140) = 0.994, p = .413, period derived aperiodic exponents.

**Figure 6.**
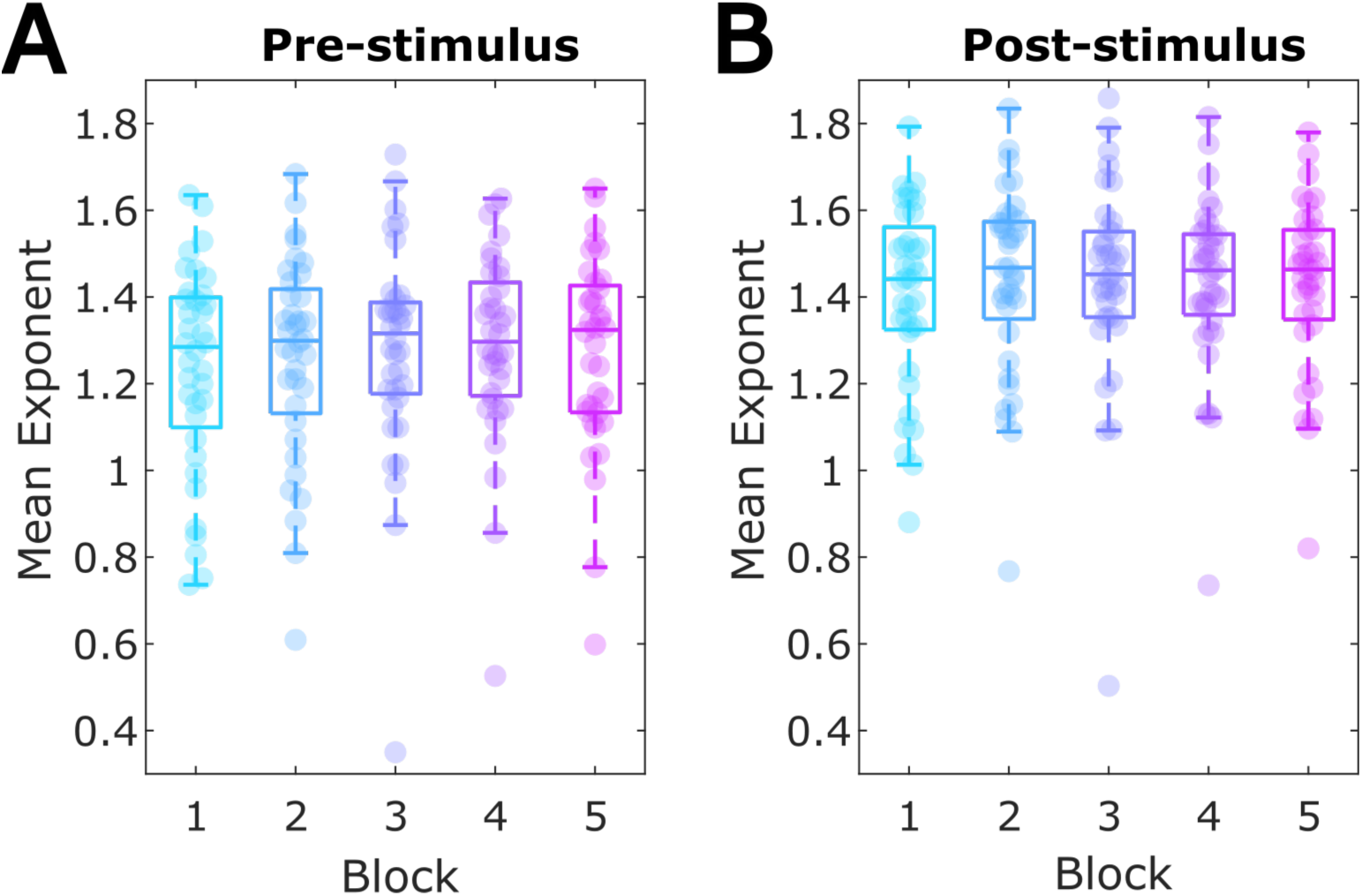
Aperiodic exponent across task blocks. (A) The mean aperiodic exponent for each block estimated for full-scalp averaged FFT spectra in the pre-stimulus period. (B) Post-stimulus exponent over blocks.

Taken together, this indicates that the non-stationarities in the alpha band across the experimental session are driven by oscillatory changes, whilst we found no evidence for time-on-task effects on the broadband aperiodic activity.

#### Time-on-task confounds relationships between alpha activity and reaction time, but not confidence

##### Time-frequency analysis

To investigate the unique relationships between EEG power and both time-on-task and behaviour with added temporal and spatial resolution, we performed a single-trial multiple regression analysis on time-frequency power data spanning -1s to 1s relative to stimulus onset.

We first tested the unique relationships between EEG power and accuracy, RTs and confidence without including time-on-task in the model. As Figure 7A shows, accuracy was unrelated to spectral power in both pre- and post-stimulus windows. Figure 7B shows RTs were significantly negatively linked to both pre- and post-stimulus alpha power, with a t_sum_ = - 48,922 and p_clust_ = .0085. Additionally, a significant positive cluster linking higher RTs to higher pre- and post-stimulus beta and low power in the gamma frequency range was also found, t_sum_ = 31,563 and p_clust_ = .0215. Higher confidence was inversely related to post-stimulus alpha and beta power (negative cluster t_sum_ = -21,819, p_clust_ = .0025) and positively related to low frequency post-stimulus power (positive cluster t_sum_ = 12,942, p_clust_ = .0135) (Figure 7C). All of the above clusters included all electrodes.

**Figure 7.**
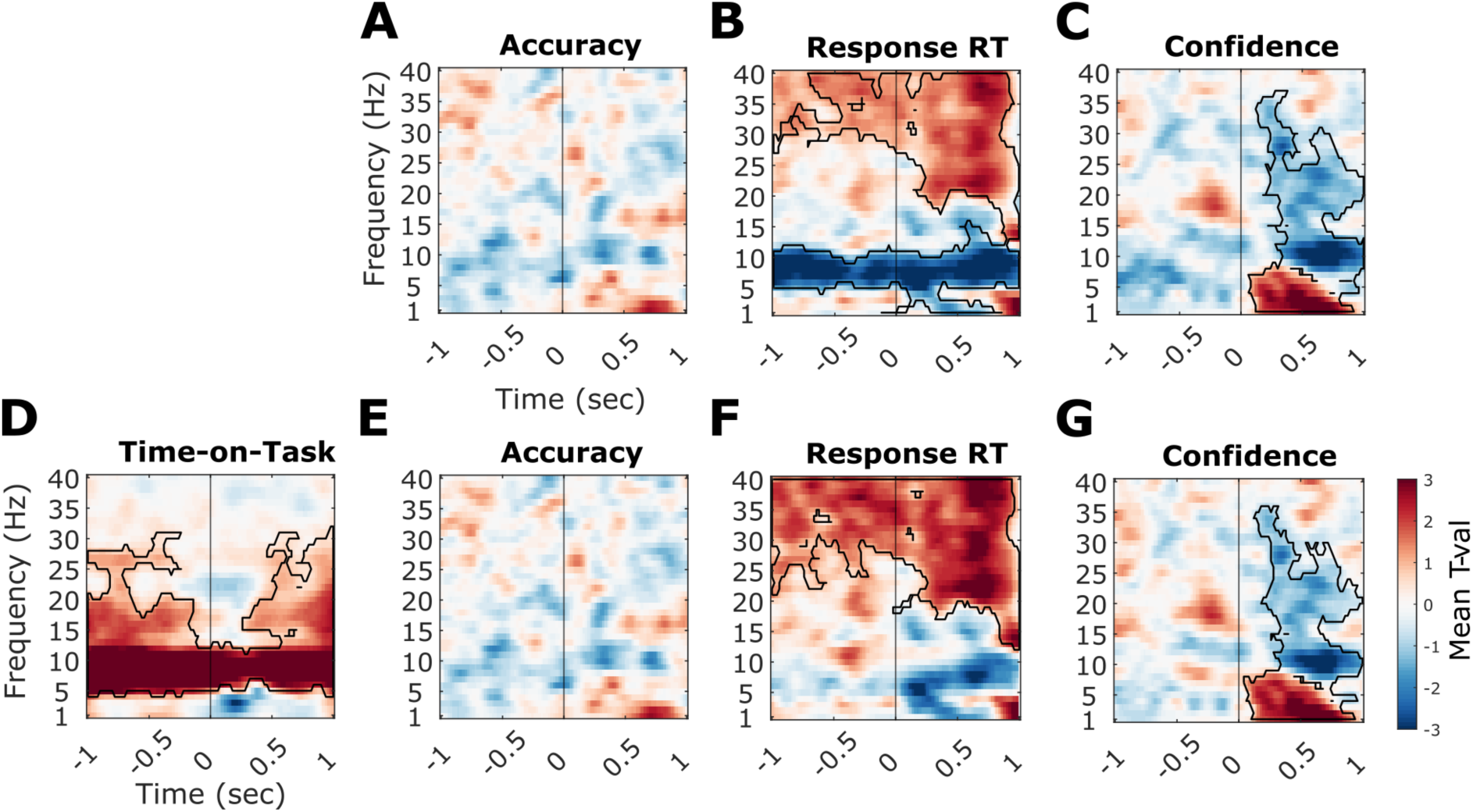
Unique relationships between EEG power and behaviour before and after including time-on-task. The time-frequency plots illustrate the mean t-values (two-tailed dependent samples t-tests of single-trial regression coefficients against zero) across all 32 electrodes that denote the strength and direction of the relationship between each behavioural measure/trial-order and spectral power. Significant clusters (p <.025) are outlined in black. The vertical line denotes stimulus onset. **A-C** Shows the unique relationships between accuracy (A), RTs (B), and confidence (C) without controlling for time-on-task. **D-G** Plot these relationships after time-on-task was controlled for. The unique effects of (D) time-on-task, (E) accuracy, (F) RTs, and (G) confidence.

To test whether any of these effects were independent from non-stationarities in the time-frequency power across the experiment, we re-ran the regression models with trial order added as another predictor. In line with the spectral analyses above, we found that time-frequency resolved power increased with time-on-task in the alpha and beta bands, with this relationship spanning across the entire pre- and post-stimulus windows in the alpha band, and being disrupted around stimulus onset in the lower beta band (Figure 7D). This effect was significant even after controlling for effects of the behavioural measures, t_sum_ = 117,107 and p_clust_ = .002. Figure 7E confirms that no significant accuracy effects were present. As Figure 7F illustrates, the reaction time effects on alpha and lower beta power (in both the pre- and post-stimulus period) disappeared after the time-on-task related variance was accounted for (max negative cluster p_clust_ = .028). However, RTs correlated with higher frequency activity (higher beta and low gamma power) in both pre- and post-stimulus periods after controlling for time on task (t_sum_ = 69,595, p_clust_ = .001) (Figure 7F). In contrast, confidence remained a significant predictor of post-stimulus alpha/beta desynchronisation and low frequency power even after time-on-task was accounted for (negative cluster t_sum_ = -17,455, p_clust_ = .0025; positive cluster t_sum_ = 14,377, p_clust_ = .006; Figure 7G), suggesting these effects are more likely due to trial-by-trial variability in spectral power rather than due to monotonous time-on-task effects. All reported clusters included all 32 electrodes.

Overall, we find that alpha power increases with time-on-task and that this effect is sustained throughout the whole pre- and post-stimulus (-1 to 1s) periods. Similarly, beta power increases with time-on-task in pre- and post-stimulus windows, although this effect is interrupted by the stimulus onset. Moreover, both these effects showed a relationship with RTs. Importantly, these effects dissociated from the relationships between confidence ratings and spectral power regarding frequencies, timing, sign, and dependence on time on task.

##### Aperiodic corrected analysis

We additionally sought to examine whether the identified time-on-task effects on purely periodic alpha activity (both frequency and power) confounded or explained relationships with behavioural performance. The behavioural measures were single-trial accuracy, reaction times, and confidence ratings, while trial stimulus contrast was controlled for in the regression models.

First, we tested the unique relationships between behaviour and both peak alpha frequency and power without accounting for time-on-task. To do so, we performed single-trial multiple regressions, on trials where an alpha peak was identified in the EEG data, separately for each participant. We then compared the resulting regression coefficients against zero to test for systematic group-level effects. As the pink bars in Figure 8 show, we found that in the pre-stimulus period (Fig. 8A-B), RTs were significantly positively related to alpha peak frequency (t=4.34, p<.001) and negatively related to alpha peak power (t=-3.07, p = .011). In contrast, neither confidence nor accuracy showed significant relationships with alpha activity in the pre-stimulus window (all p’s > .05). In the post-stimulus data (Fig. 8D-E), RTs showed a significant positive relationship with alpha peak frequency (t=3.43, p = .008), whilst the other behavioural measures were unrelated to alpha frequency. Post-stimulus alpha peak power, on the other hand, was significantly negatively linked to single trial confidence ratings (t=-5.25, p <.001) and RTs (t=-3.21, p= .011), but unrelated to accuracy (p = .198).

**Figure 8.**
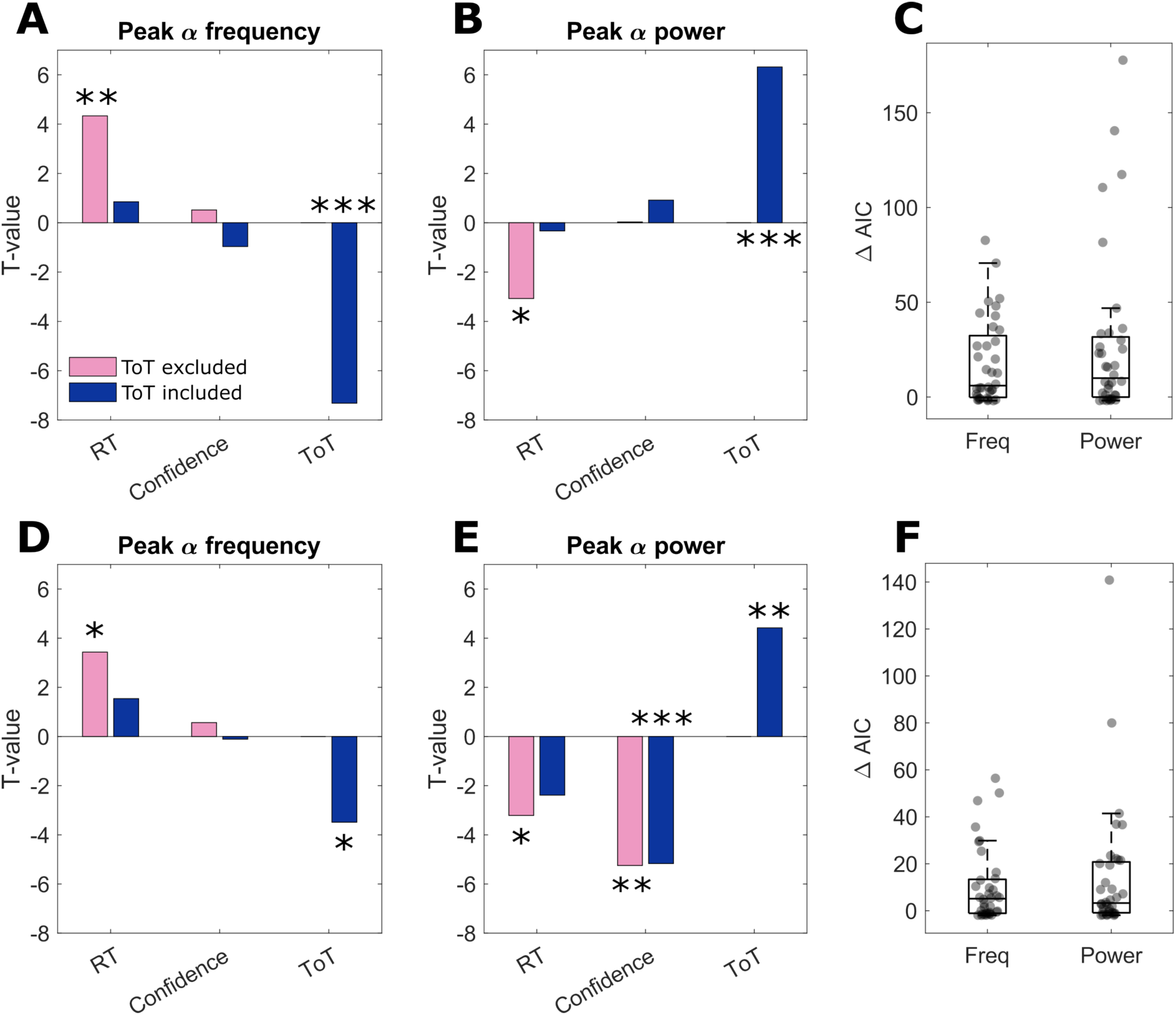
**A-B** Pre-stimulus and **D-E** post-stimulus comparisons of results of multiple regression testing (with group-level t-tests) for unique effects of behaviour on EEG with (in blue) and without (in pink) controlling for time-on-task (ToT). * *p* <.05; ** *p* <.001, *** *p* <.0001 (after FDR multiple comparisons corrections (Benjamini & Hochberg, 1995)). **C** (pre-stimulus) and **F** (post-stimulus) show the difference in AIC between the models without and with ToT – positive ΔAIC denote better model fits with ToT included.

Hence, alpha power and frequency, in both pre- and post-stimulus periods, were significantly correlated with a time-varying behavioural measure, namely response RTs. However, given the time-on-task effects on alpha activity and response speed these relationships may be confounded by trial order.

We therefore repeated the regression analyses whilst also including trial order as an additional predictor in the models. The blue bars in Figure 8 show the results of the group-level t-tests of regression coefficients against zero. We found that when time-on-task was controlled for, neither pre-stimulus alpha peak frequency nor peak power predicted RTs (p = .572, p = .874). However, time-on-task itself remained a significant unique predictor of both pre-stimulus alpha frequency (t=-7.32, p <.0001) and power (t=6.32, p <.0001) even after controlling for the effects of behavioural measures. No other significant relationships were found with confidence or accuracy in the pre-stimulus period (all p’s > .05).

Like the pre-stimulus effects, the previously significant relationships between post-stimulus alpha frequency (p = .264) and power (p = .065) and RTs did not reach significance after time-on-task effects were accounted for. Additionally, post-stimulus alpha frequency decreased significantly with time-on-task (t=-3.49, p = .005) and alpha power increased (t=4.42, p = .0005) even when all behavioural measures were controlled for. Whilst post-stimulus alpha frequency was unrelated to any behavioural measures (p’s > .05), alpha power remained inversely related to confidence after accounting for time-on-task effects (t=-5.17, p <.0001).

To formally test whether the inclusion of time-on-task significantly improved the models, we calculated the difference in AIC values for the model fits with and without time-on-task included (Figure 8C&F). We found that the regression models including time-on-task had better model fit than the models without time-on-task. This result was found for both alpha power and frequency in the pre- and post-stimulus time windows (all p’s <.005; in t-tests comparing ΔAIC against 0 (i.e., no improvement in fit)).

Together these results suggest that time-on-task effects fully accounted for relationships between EEG alpha activity and RTs, while post-stimulus relationships between EEG alpha power and confidence occurred independently of time-on-task.

## Discussion

We examined how time-on-task influences both periodic and aperiodic EEG activity across the course of a typical experimental session, and whether time-on-task trends potentially confound apparent relationships between EEG and behaviour. During performance of a visual discrimination task, neural time-on-task trends were found in the alpha (8-12Hz), and beta bands (13-20Hz). Specifically, alpha (and beta) power systematically increased over the course of task performance, whereas alpha frequency systematically decreased. The effects represent changes in oscillatory rather than aperiodic activity: aperiodic activity did not vary systematically across the experiment. Importantly, time-on-task confounded apparent pre- and post-stimulus relationships between single-trial alpha activity and reaction times which were no longer significant after time-on-task was accounted for. These results establish the oscillatory nature of long-term EEG non-stationarities in the alpha band and emphasise the need to account for time-on-task when testing oscillatory brain-behaviour relationships. We argue that controlling for time-on-task ultimately allows for a better understanding of the functional roles of oscillations in cognition as it may remove potentially spurious correlations and allow to tease apart different oscillatory contributors to behavioural variability.

The increases in alpha and beta power across task blocks are in line with previous studies that found time-on-task spectral changes (Benwell et al., 2019; Boksem et al., 2005; Cajochen et al., 1995; Sadaghiani et al., 2010; Stoll et al., 2016). These power increases across longer timescales have previously been linked to fatigue and lapses in sustained attention (Boksem et al., 2005; Cajochen et al., 1995). Additionally, Benwell et al. (2019) showed that alpha band non-stationarities likely arise from different underlying mechanisms, with potentially separable functions, as some alpha components showed changes in power, frequency, or combination of both in their study. Here, by examining the time-on-task effects across a wider frequency spectrum, we confirm the previous findings and show that alpha band activity experiences a relatively steep increase, accompanied by a shallower increase in the beta band (see Figure 4). Whilst our task did not elicit a decline in performance across the experimental session (i.e., more errors/longer response times) with time-on-task on the group level, it is possible that the alpha power increase may at least partially reflect increasing fatigue within-participants across the experimental session.

The increases in beta power are also somewhat in line with Stoll et al.’s (2016) findings of time-on-task effects on beta activity in monkeys, who argued that this trend may reflect increasing attentional effort required to maintain cognitive performance. It remains unclear whether the beta effect is a distinct phenomenon or a consequence of alpha harmonics that carry the alpha power trend into the beta band. However, the differences in slope, with alpha power showing a steeper increase, and different topographical representations may suggest at least partially independent processes.

Our results show that systematic non-stationarities in alpha band activity are driven by oscillatory changes and not due to alterations in the aperiodic signal. This is important given that previous time-on-task studies (Benwell et al., 2019; Boksem et al., 2005; Cajochen et al., 1995; Sadaghiani et al., 2010) did not account for the aperiodic exponent, which has been shown to confound spectral results regarding EEG changes with aging (Merkin et al., 2021; Tröndle et al., 2022), cognitive performance (Ouyang et al., 2020), and disease states (Johnston et al., 2023; Pani et al., 2022; Robertson et al., 2019). We found no significant changes in the aperiodic exponent across blocks. On the physiological level, this implies that the EEG non-stationarities are unlikely to be due to changes in asynchronous neural spiking (Manning et al., 2009) or alterations in the balance between excitatory and inhibitory synaptic currents (Gao et al., 2017), which have both been linked to the aperiodic component. Instead, our results suggest that time-on-task related changes arise from alteration in neural synchrony in widespread cortical networks which are thought to give rise to oscillations (Buzsáki et al., 2013; Buzsáki & Draguhn, 2004). The concurrent increases in alpha power and decreases in alpha frequency have previously been attributed to a single process and proposed to reflect fluctuations in the background excitability (Himmelstoss et al., 2015). However, Benwell et al (2019) showed that the changes in alpha power and frequency arise from several distinct generating sources with separable spatial and spectral characteristics. Our results may therefore reflect changes in one or more oscillators that vary over the course of an experimental session. Considering the numerous functional roles ascribed to alpha activity, including task-related functional inhibition (Foxe & Snyder, 2011), information integration and transfer (Griffiths et al., 2019; Jensen & Mazaheri, 2010), and sensory sampling (Samaha & Postle, 2015), the functional relevance of the time-varying alpha activity remains to be determined.

Our results highlight the importance of considering time-on-task effects when interpreting brain-behaviour relationships. In addition to non-stationarities in the EEG signal, we found that participants’ accuracy improved, and response times became faster over the initial blocks of the experiment, which could be due to practice effects, whereas confidence remained unchanged. Previous studies have found that links between some behavioural measures and EEG activity can be at least partially attributed to deterministic sources of variance, like time-on-task related shifts in both EEG and behaviour (Benwell et al., 2018; Bompas et al., 2015; Schaworonkow et al., 2015). Our results are in line with this, as we found that time-on-task effects confounded relationships between alpha activity and reaction times. When time-on-task has not been accounted for, numerous studies have found links between reaction times and alpha power (Dockree et al., 2007; Kelly et al., 2009; Roberts et al., 2014), as well as frequency (Jin et al., 2006), across a variety of tasks. Accordingly, we also found that higher alpha power was linked to faster RTs, whereas higher alpha frequency was related to faster RTs across both pre- and post-stimulus windows. However, adding time-on-task as a regressor in our model explained away the direct relationship between alpha power and response speed completely.

This was true in both spatially resolved time-frequency analyses as well as in scalp-averaged analyses in which only purely oscillatory activity was included. This finding highlights how correlations between two measures affected by time-on-task can arise spuriously and can be misinterpreted as a direct functional relationship. Hence, we strongly advise including time-on-task as a default control parameter when studying oscillation-behaviour links.

In addition to time-on-task related EEG effects, we also found that alpha and beta desynchronisation uniquely predicted single-trial decision confidence independently of accuracy, RTs, stimulus contrast, and trial-order. This is in line with previous studies that showed post-stimulus alpha/beta power correlates with subjective judgements like confidence and perceptual awareness (Benwell et al., 2022; Faivre et al., 2018; Kopčanová et al., 2023) and is unrelated to accuracy (Benwell et al., 2022; Kopčanová et al., 2023). Given the independence of this effect from RTs, it is unlikely it can solely be attributed to motor preparation (Faivre et al., 2018). Instead, it may reflect fluctuations in excitability and global arousal (Dahl et al., 2022; Kosciessa et al., 2021) that uniquely affect confidence and not accuracy (Allen et al., 2016). Importantly, the finding that this effect persists after controlling for time-on-task, i.e., is not a spurious consequence of both alpha/beta and confidence changing over time, highlights that this approach can help separating spurious from genuine effects of neural oscillations on cognitive function. It may also indicate the existence of different alpha-generating processes that tie in with distinct physiological and cognitive processes.

In addition to time-on-task independent alpha/beta effects, we also found that low frequency (delta/theta) power predicted single trial confidence judgements independently of time-on-task and other behavioural measures. It is likely that the positive correlations between delta/theta power and confidence reflect the previously found relationships between slow ERPs (e.g., P300) and accumulating decision evidence as well as confidence judgements (Feuerriegel et al., 2022; Gherman & Philiastides, 2018; Herding et al., 2019; Kelly & O’Connell, 2013; Kopčanová et al., 2023; Trajkovic et al., 2024).

Finally, we also found slower reaction times were linked to increases in high beta/low gamma power even after time-on-task was controlled for. Increased pre-stimulus gamma power has previously been linked to slower RTs (Reinhart et al., 2011), whereas others have found a negative relationship between gamma and RTs (Hoogenboom et al., 2010). Whilst our results are more in line with Reinhart et al.’s (2011) who attributed it to a state of preparedness that leads to more complete stimulus processing and slower motor responses, we cannot exclude the possibility that this effect is driven by muscle-related activity in the current study. Nevertheless, these effects overall demonstrate that EEG-behaviour relationships outside of the alpha/beta band are unlikely to reflect deterministic variance due to time-on-task.

## Conclusion

We found strong systematic time-on-task trends in the alpha band, with weaker effects present in the beta band. These trends were absent in aperiodic activity, corroborating earlier findings of long-term changes in the activity of cortical oscillators during sustained task performance. Our findings also emphasise the importance of accounting for time-on-task when investigating brain-behaviour relationships, as monotonous changes in both EEG and behavioural measures may drive correlations that can be misinterpreted. Finally, we provide evidence that time-varying performance-related alpha/beta effects can be dissociated depending on whether they can be accounted for by time on task, here: response speed, or whether they persist after controlling for it, here: decision confidence.

## Acknowledgments

MK is funded by an SGSSS (UKRI ESRC: ES/P000681/1) stipend awarded to authors CK, CB and GT. All authors are members of the Scottish-EU Critical Oscillations Network (SCONe), funded by the Royal Society of Edinburgh (RSE Saltire Facilitation Network Award to CK, Reference Number 1963). The authors would also like to thank Efstratios Koukouvinis for his help with data collection.

