## Supplementary for "Characterising time-on-task effects on oscillatory and aperiodic EEG components and their co-variation with visual task performance"

| # | **Author** | **Affiliation** | **ORCID** | **Social Media** |
|---|---|---|---|---|
| 1 | Martina Kopčanová | 1 | 0009-0004-0300-3343 | |
| 2 | Gregor Thut | 2 | 0000-0003-1313-4262 | |
| 3 | Christopher SY Benwell | 1 | 0000-0002-4157-4049 | |
| 4 | Christian Keitel | 1 | 0000-0003-2597-5499 | @ckeitelsci.bluesky.social (BlueSky) |


### Keywords

EEG, neural oscillations, alpha, time on task, reaction time

**Table S1** Results of one-way ANOVAs with evidence discriminability as IV and RT as DV tested separately at each block.

| Block | F(4,140) | $p_{gg}$ |
|---|---|---|
| 1 | **19.699** | **<.0001** [a] |
| 2 | **8.170** | **.0025** [b] |
| 3 | **13.783** | **<.0001** [c] |
| 4 | 3.187 | .0513 |
| 5 | **10.709** | **<.0001** [d] |

The follow up multiple comparisons were significant p < .05 between the following blocks:
[a] block 1 versus 2, 3, 4, and 5; and block 2 vs 3, 4, and 5.
[b] block 1 vs 3, 4, and 5.
[c] block 1 vs 3, 4, and 5; block 2 vs 3, 4, and 5.
[d] block 1 vs 2, 3, 4, and 5; and block 2 vs 5.

**Table S2** Results of one-way ANOVAs with block as IV and RT as DV tested separately at each evidence discriminability level.

| Difficulty level | F(4,140) | $p_{gg}$ |
|---|---|---|
| **Hard to Easy** | | |
| 1 | **22.339** | **<.0001** [a] |
| 2 | **24.063** | **<.0001** [b] |
| 3 | **26.085** | **<.0001** [c] |
| 4 | **20.902** | **<.0001** [d] |
| 5 | **10.629** | **<.0001** [e] |

The follow up multiple comparisons were significant p < ,05 between the following blocks:
[a, b, c, d, e] block 1 versus 2, 3, 4, and 5.

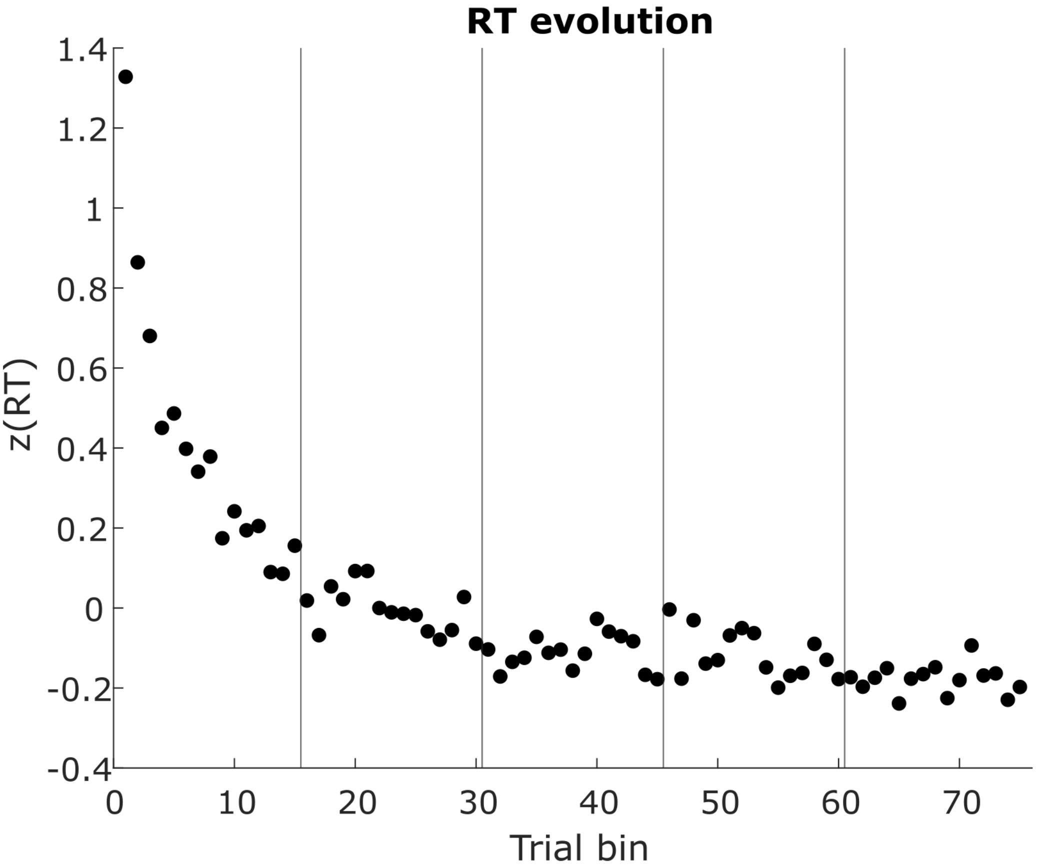

**Figure S1:** Response reaction times evolution plot. Mean (z-scored) RTs per bin of 10-12 trials in each experimental block were calculated, similar to alpha and beta power in Figure 4 in the main text.
